# Influence of oceanography and geographic distance on genetic structure: how varying the sampled domain influences conclusions in *Laminaria digitata*

**DOI:** 10.1101/2023.05.11.540379

**Authors:** L. Fouqueau, L. Reynes, F. Tempera, T. Bajjouk, A. Blanfuné, C. Chevalier, M. Laurans, S. Mauger, M. Sourisseau, J. Assis, L. Lévêque, M. Valero

## Abstract

Understanding the environmental processes shaping connectivity can greatly improve management and conservation actions which are essential in the trailing edge of species’ distributions. In this study, we used a dataset built from 32 populations situated in the southern limit of the kelp species *Laminaria digitata*. By extracting data from 11 microsatellite markers, our aim was to (1) refine the analyses of population structure, (2) compare connectivity patterns and genetic diversity between island and mainland populations and (3) evaluate the influence of sampling year, hydrodynamic processes, habitat discontinuity, spatial distance and sea surface temperature on the genetic structure using a distance-based redundancy analysis (db-RDA). Analyses of population structure enabled to identify well connected populations associated to high genetic diversity, and others which appeared genetically isolated from neighboring populations and showing signs of genetic erosion verifying contrasting ecological (and demographic) status in Brittany and the English Channel. By performing db-RDA analyses on various sampling sizes, geographic distance appeared as the dominant factor influencing connectivity between populations separated by great distances, while hydrodynamic processes were the main factor at smaller scale. Finally, Lagrangian simulations enabled to study the directionality of gene flow which has implications on source-sink dynamics. Overall, our results have important significance in regard to the management of kelp populations facing pressures both from global warming and their exploitation for commercial use.

## 1. INTRODUCTION

Understanding connectivity patterns in endangered species living in fragmented habitats is fundamental to assess their vulnerability to global changes and define conservation measures at appropriate scales (*e.g.*, Manel et al. 2019). Such knowledge is particularly relevant at the trailing edge of species’ distribution, where global warming effects interfere with the strong genetic and demographic drifts (Nadeau & Urban 2019). Investigating connectivity can additionally pinpoint populations that are isolated from gene flow, which can impede adaptation and ultimately lead to extinction (Molofsky & Ferdy 2005, Allendorf & Luikart 2007). A deeper understanding of populations’ connectivity can therefore advance conservation action aiming to preserve species’ adaptive potential via measures that prevent the loss of genetic diversity.

Most marine organisms disperse through planktonic propagules (*e.g*., spores, eggs, larvae) which are deemed unable of countering hydrodynamic forces (Alberto et al. 2010), although life history traits and behavior seem to influence dispersal kernels to a certain extent (Sanford & Kelly 2011). Therefore, since connectivity among marine species’ has begun to receive some attention, the influence of oceanographic processes on contemporary genetic structures have quickly been proven by theoretical and empirical works (*e.g.*, Gilg & Hilbish 2003; Treml et al. 2008; Mitarai et al. 2009; Alberto et al. 2010; White et al. 2010; D’Aloia et al. 2014, Thibaut et al. 2016). For instance, hydrodynamic processes have been associated with larval retention and asymmetrical gene flow (Gaylord & Gaines 2000), both of which inevitably affect metapopulation dynamics. Additionally, White et al. (2010) demonstrated that ocean currents enable exchanges between island populations, which were historically considered genetically isolated due to geographic isolation and high endemism (Cowen et al. 2000). However, to this date, a formal comparison of the connectivity between island and coastal populations is still poorly explored for marine species compared to terrestrial ones (Bell 2008). Although oceanic currents are fundamental in shaping the genetic structure among marine species, other features need to be considered in order to deepen our understanding of connectivity (Hu et al. 2020). Finally, processes influencing the genetic structure are expected to differ over different geographical extents, emphasizing the importance of considering different spatial scales when exploring their relative contribution (Jombart et al. 2009, Dalongeville et al. 2018).

Seascape genetics aims to investigate which and how marine environmental features influence the spatial distribution of genetic variation at neutral loci or the ones presumably under selection. Among the tested features, water depth (*e.g.*, Engel et al. 2004; Hickey et al. 2009; Krueger-Hadfield et al. 2013), habitat discontinuity (*e.g.*, Johansson et al. 2008; Fraser et al. 2009; Alberto et al. 2010, 2011; D’Aloia et al. 2014; Durrant et al. 2018), sea surface temperature (termed SST hereafter, *e.g*., Johansson et al. 2015; Benestan et al. 2016; Guzinski et al. 2020) and salinity (*e.g.*, Gaggiotti et al. 2009; Sjöqvist et al. 2015) have been shown to explain, separately or jointly, a large part of the variation in genetic structure. Most of these studies have relied on linear regressions and Mantel tests to show the influence of environmental factors (*e.g.*, White et al. 2010, Alberto et al. 2011, and see Benestan et al. 2016 for further references) but these statistical frameworks have since been shown to be inappropriate. While linear regressions appear unsuited due to the violation of the assumption of independency between *F_ST_* values (Benestan et al., 2016; Boldina & Beninger, 2016), Mantel tests lead to a large decrease in statistical power and can provide erroneous conclusions (Legendre & Fortin, 2010). To overcome these limitations and adequately tackle distance variables, distance-based redundancy analyses (db-RDA) have been put forward by Legendre & Anderson (1999). Db-RDA is a direct extension of multiple regression and models linear combinations of explanatory variables, enabling a more accurate evaluation of the relative contribution of each explanatory variable. In addition to these statistical advances, the improvement of hydrodynamic models and the development of Moran’s and asymmetric eigenvector maps (Dray et al. 2006, Blanchet et al. 2008) allowed a better apportioning of the contribution of geographic distance and oceanographic features (*e.g.*, Benestan et al. 2016, 2021; Xuereb et al. 2018; Reynes et al. 2021).

Kelp forests are a dominant feature along many temperate to boreal rocky shores and play a foundation role for numerous species by providing them substrate, shelter or food (Teagle et al. 2017, Jayathilake & Costello 2020, Coleman & Veenhof 2021). The decline they have shown in some biogeographic regions has justified their recent inclusion in the OSPAR list of threatened and declining habitats (see de Bettignies et al. 2021). SST is considered as the major abiotic factor shaping kelp species’ range distribution as denoted by the cold-water niches they tend to be associated with (Lüning 1990, Bartsch et al. 2008). Over the past decade, various studies have shown the harmful consequences of the gradual increase in SST on kelp forests, notably as a result of ever more frequent and intense marine heatwaves (Filbee-Dexter et al. 2016, Wernberg et al. 2016, Starko et al. 2019, Cavanaugh et al. 2019, Rogers-Bennett & Catton 2019, Coleman et al. 2020), which especially affect warm-edge populations (Fernández 2011; Arafeh-Dalmau et al. 2019; Starko et al. 2019; Filbee-Dexter et al. 2020 but see Klingbeil et al. 2022). In this context, clarifying connectivity in the warm edge could be particularly valuable to gauge the resilience of marginal populations. For instance, detecting dispersal from the range center towards the margin would be a positive sign of adaptive potential, especially if marginal and core populations are submitted to similar environmental conditions (Bridle et al. 2009, DuBois et al. 2022).

*Laminaria digitata* (Hudson) J.V. Lamouroux is a boreal kelp species with an amphi-Atlantic distribution. Its geographic range in the northeast Atlantic extends from temperate southern Brittany (47°N, France) to the arctic Spitsbergen archipelago (79°N, Norway) (Kain 1979, Lüning 1990, Araújo et al. 2016). Along the shorelines, *L. digitata* generally occurs within a narrow band (ca. 5 to 10 m wide) spanning the lower intertidal and upper subtidal zones (Robuchon et al. 2014). This species thus presents an interesting case to test the effect of coastal oceanographic currents on genetic structure, especially along the rugged coast of Brittany, where hydrodynamics is highly complex due to numerous mesoscale features (*e.g.*, fronts, upwelling) and macrotidal ranges (Salomon & Breton 1993, Billot et al. 2003, Ayata et al. 2010, Nicolle et al. 2017). These studies have also identified isolated groups of populations, although the causal environmental factors have not been formally identified. Additionally, annual mean SST varies at a fine scale across Brittany, with the western and north-western sectors being cooler and currently less affected by climate change than Southern and North-Eastern Brittany (Gallon et al. 2014). As suggested by Liesner et al. (2020), this mosaic of SST conditions might have driven the distinct thermal adaptations observed between populations from northern and southern Brittany. Finally, by using a hierarchical sampling scheme, previous analyses on population structure at the scale of Brittany have revealed significant genetic differentiation at both small (< 1 km) and large scale (> 10-50 km) relative to the region (Billot et al., 2003; Robuchon et al., 2014).

In this study, we extended the sampling design of previous microsatellite analyses (Billot et al. 2003, Valero et al. 2011, Couceiro et al. 2013, Robuchon et al. 2014) by using a dataset built from 32 populations ranging from the southern range limit of *L. digitata* to the northernmost population found on the French coast, situated in the Strait of Dover. Using this dataset, we explored the respective and combined effects of sampling year, hydrodynamic processes, habitat discontinuity, spatial distance and SST on the genetic structure at different domain sizes. We specifically aimed to (1) refine the analyses of population structure for this species and identify isolated populations, (2) test whether island populations are disconnected from mainland populations, and thus show signs of genetic erosion and (3) evaluate the relative importance of the environmental factors on the genetic structure at different sampling domain sizes using db-RDA. In our study, we defined “island” populations as those corresponding to on-shelf islands since our goal was to compare connectivity among coastal populations, among islands or between coastline and island populations for equivalent geographical distances.

## 2. MATERIAL AND METHODS

### 2.1. Study area

Brittany is the northwestern-most region of mainland France (NE Atlantic). Its 2,860 km-length coastline encompasses a major biogeographical transition zone between cold- and warm-temperate waters (Spalding et al. 2007). The northern shores, which border the English Channel, are characterized by well-mixed waters produced by a macrotidal regime that intensifies eastwards. Contrastingly, waters in southern Brittany are seasonally to permanently stratified and show larger temperature fluctuations throughout the year (*e.g*., Blauw et al., 2019). Brittany’s western-most sector is subject to both the macrotidal regime and the full impact of Atlantic storms, which collectively maintain well-mixed conditions throughout the year. Various boreal-affinity species benefit from this nutrient-rich and thermally-buffered environment, including various kelp species that form some of Europe’s most important marine forests along Brittany’s rocky shores.

### 2.2. Samples

Populations of *L. digitata* were sampled between 2005 to 2015 in 32 sites ranging from the southern limit of its North-East Atlantic distribution (47° latitude) up to the Strait of Dover, at the entrance of the North Sea (50° latitude, Table 1, Figure 1). The population at the Strait of Dover is known to be isolated from other populations, with the closest population along the French coast being the one from Étretat, which is still situated at *circa* 200 km south (see Figure 2C of Araújo et al. 2016) and for which we did not have any sample. Among our populations, 10 were located around islands while 22 were placed on the mainland coast (Table 1, Figure 1). Some of these populations were sampled and genotyped for previous studies as indicated in Table 1. At each site, blade tissue was collected from 30 to 50 randomly selected sporophytes. Tissue samples were then wiped, cleaned from epiphytes and stored in silica-gel crystals until DNA extraction.

**Figure 1.**
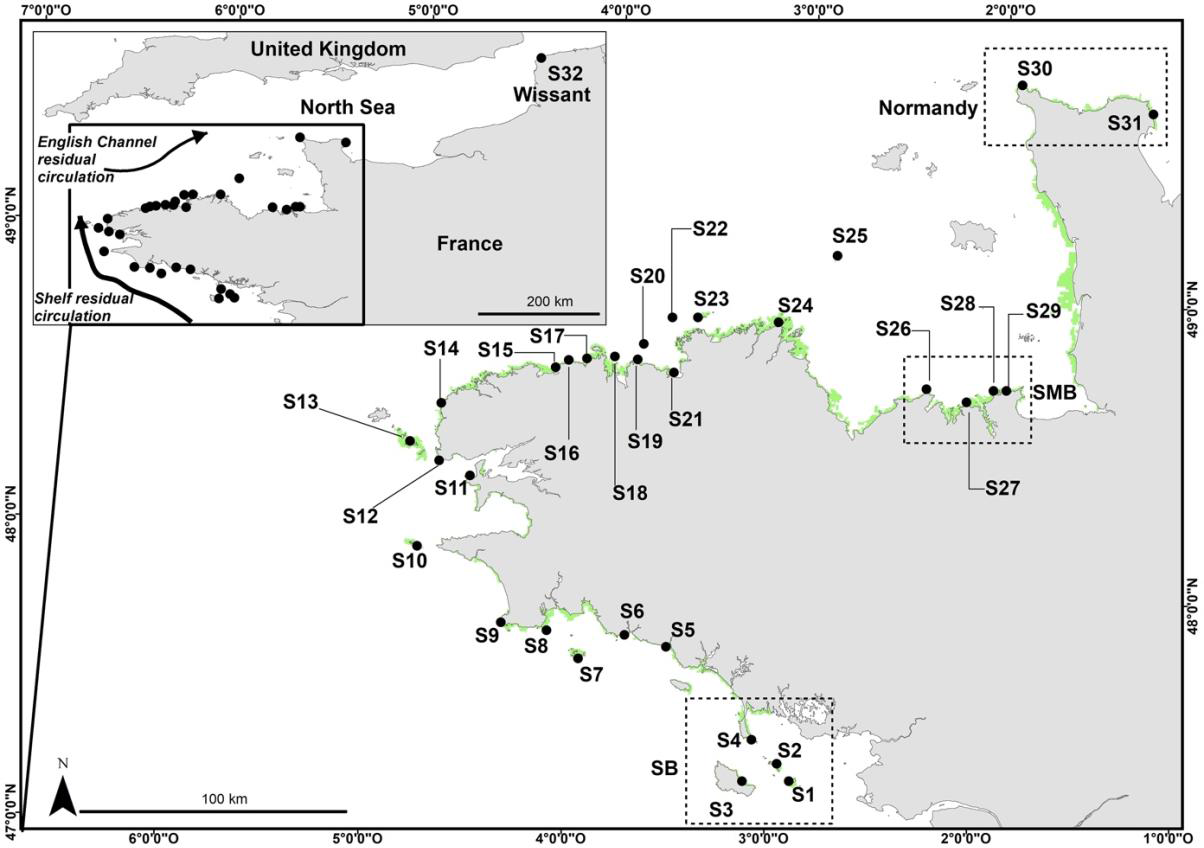
Geographic position of the sampled populations, the Northernmost site (S32) is indicated in the inset. Population numbers correspond to the ones indicated in Table 1 and the area illustrated in green corresponds to rocky substrata above 5m depth obtained from IFREMER in May 2019. The information on the spatial distribution of bedrock comes from the combination of several sources as specified in the Material and Method section. The inset illustrates the general circulation in the Bay of Biscay and in the English Channel, drawn according to Ayata et al. (2010).

**Figure 2.**
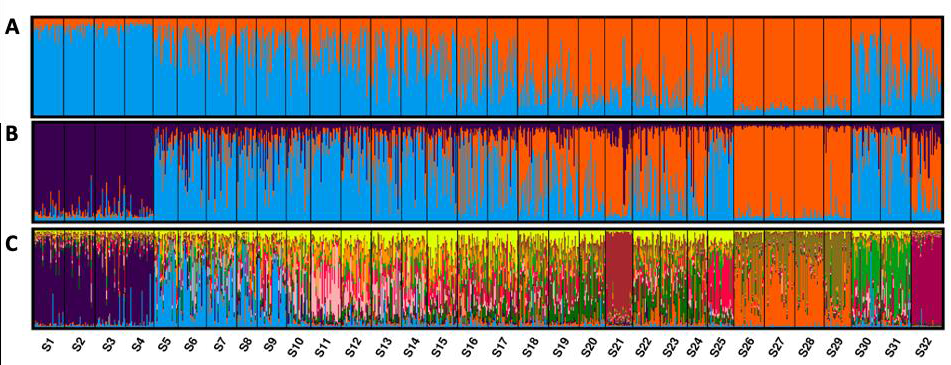
STRUCTURE barplot obtained for K = 2 (**A**), K = 3 (**B**) and K = 12 (**C**) which appeared the best number of clusters according to both ΔK and log Pr(X|K) methods. Individuals corresponding to vertical bars are assigned to each cluster with a certain probability. The numbers above the Figure C correspond to population number as indicated in Table and Figure 1.

**Table 1.** Information on the sampled populations: populations’ number as indicated in Figure 1; geographic locations (Latitude, Longitude); name of the sampling locality; year of sampling; position in regard to the coastline; number of genotyped individuals and source of the data.

| N° | Lat. | Long. | Locality | Year | Position | Nb. Ind. | Source |
| --- | --- | --- | --- | --- | --- | --- | --- |
| S1 | 47.339 | -2.892 | Hoedic | 2011 | Island | 30 | Robuchon et al. (2014) |
| S2 | 47.394 | -2.958 | Houat | 2011 | Island | 30 | Robuchon et al. (2014) |
| S3 | 47.328 | -3.124 | Belle-île | 2006 | Island | 30 | This study |
| S4 | 47.470 | -3.091 | Quiberon | 2008 | Coastline | 28 | This study |
| S5 | 47.762 | -3.549 | Lorient | 2006 | Coastline | 24 | Valero et al. (2011) |
| S6 | 47.791 | -3.761 | Pont Aven | 2006 | Coastline | 28 | Valero et al. (2011) |
| S7 | 47.700 | -3.983 | Glénan | 2008 | Island | 30 | This study |
| S8 | 47.786 | -4.152 | Loctudy | 2006 | Coastline | 29 | This study |
| S9 | 47.799 | -4.383 | Penmarc'h | 2008 | Coastline | 20 | Valero et al. (2011) |
| S10 | 48.032 | -4.835 | Ile de Sein | 2009 | Island | 24 | This study |
| S11 | 48.284 | -4.601 | Crozon | 2011 | Coastline | 30 | This study |
| S12 | 48.326 | -4.764 | Le Conquet | 2011 | Coastline | 30 | Robuchon et al. (2014) |
| S13 | 48.381 | -4.919 | Le Conquet | 2011 | Island | 30 | Robuchon et al. (2014) |
| S14 | 48.519 | -4.779 | Porspoder | 2011 | Coastline | 30 | Robuchon et al. (2014) |
| S15 | 48.673 | -4.216 | Plouescat | 2005 | Coastline | 30 | Valero et al. (2011) |
| S16 | 48.700 | -4.152 | Téven-Meur | 2011 | Coastline | 30 | Robuchon et al. (2014) |
| S17 | 48.711 | -4.060 | Ile de Sieck | 2005 | Coastline | 30 | Valero et al. (2011) |
| S18 | 48.725 | -3.919 | Roscoff | 2011 | Coastline | 30 | Robuchon et al. (2014) |
| S19 | 48.721 | -3.803 | Plougasnou | 2011 | Coastline | 30 | Robuchon et al. (2014) |
| S20 | 48.775 | -3.777 | Plateau de la Méloine | 2015 | Island | 26 | This study |
| S21 | 48.687 | -3.613 | Locquirec | 2006 | Coastline | 27 | Valero et al. (2011) |
| S22 | 48.871 | -3.643 | Triagoz | 2012 | Island | 27 | This study |
| S23 | 48.878 | -3.513 | Trégastel | 2006 | Island | 27 | This study |
| S24 | 48.881 | -3.099 | Tréguier | 2006 | Coastline | 20 | This study |
| S25 | 49.118 | -2.821 | Roche Douvres | 2012 | Island | 26 | This study |
| S26 | 48.688 | -2.325 | Plévenon | 2011 | Coastline | 30 | Robuchon et al. (2014) |
| S27 | 48.651 | -2.118 | Saint-Lunaire | 2011 | Coastline | 30 | Robuchon et al. (2014) |
| S28 | 48.694 | -1.984 | Saint-Malo | 2011 | Coastline | 29 | Robuchon et al. (2014) |
| S29 | 48.697 | -1.919 | Saint-Malo | 2006 | Coastline | 27 | This study |
| S30 | 49.729 | -1.919 | Omonville | 2006 | Coastline | 29 | This study |
| S31 | 49.653 | -1.234 | Barfleur | 2006 | Coastline | 30 | This study |
| S32 | 50.912 | 1.677 | Wissant | 2006 | Coastline | 30 | This study |

### 2.3. Microsatellite genotyping

#### 2.3.1. DNA extraction

DNA was extracted from 8–12 mg of dried tissue using the NucleoSpin 96 Plant II kit (Macherey-Nagel GmbH & Co. KG) following manufacturer’s instructions. The lysis, microsatellite amplification and scoring were performed for 12 polymorphic loci following Robuchon et al. (2014). Multiplex PCRs were modified using 5X GoTaq Flexi colorless reaction buffer (Promega Corp., Madison, USA) instead of 1X and performed using a T100™ Thermal Cycler (Bio-Rad Laboratories Inc.).

#### 2.3.2. Microsatellite amplification, scoring, and correction

Among the markers used for this study, six were previously developed for *L. digitata* (Ld148, Ld158, Ld167, Ld371, Ld531, and Ld704; Billot et al. 1998) and five for *L. ochroleuca* (Lo4-24, Lo454-17, Lo454-23, Lo454-24, and Lo454-28; Coelho et al. 2014). Alleles were sized using the SM594 size standard (Mauger et al. 2012) and scored manually using GeneMapper 4.0 (Applied Biosystems). Individuals for which more than one locus did not amplify were removed from the dataset.

#### 2.3.3. Preliminary analyses

Prior to genetic analyses, a Principal Component Analysis (PCA) was ran using the “adegenet” R package (Jombart 2008) to identify potential outliers. Then the presence of null alleles was estimated on the dataset cleaned from potential outliers, using the ENA method in FreeNa (Chapuis & Estoup 2007). As we expect the frequency of null alleles to increase with mutation rate, we also tested the occurrence of null alleles at each marker by looking at the correlation between the mean number of observed alleles (*N_a_*) and the number of individuals for which there was no amplification. In order to test the independence of each marker, genotypic linkage disequilibrium was calculated for all pairs of markers among each of the populations and across populations using Genepop 4.7.5 (Rousset 2008) and the Markov chain parameters that were used are: dememorization number 1,000, number of batches 100 and number of iterations per batch 1,000.

### 2.4. Population structure

Population structure was investigated with pairwise estimates of *F_ST_* (Weir & Cockerham 1984) and their significance was computed with 1,000 permutations on FSTAT (Goudet 1999). Population structure was additionally assessed with two multivariate methods: a Principal Coordinate Analysis (PCoA) using the R package “ape” (Paradis & Schliep 2019) and STRUCTURE v2.3.4 (Pritchard et al. 2000). The PCoA represents the first step to run the multivariate statistical procedure (db-RDA, see section “Linking environmental variables to genetic structure”). STRUCTURE was run with the admixture model without prior population information and ten independent replicate runs were performed from K = 1 to 20 rather than 32 (corresponding to the number of populations) to reduce computation time. For each replicate, the burn-in was set to 100,000 and the Markov chain Monte Carlo (MCMC) iterations to 500,000 following guidance from Gilbert et al. (2012). The most likely value of K was determined using both the Evanno ΔK (Evanno et al. 2005) and the log Pr(X|K) methods (Pritchard & Wen 2003) obtained using Structure Harvester (Earl & VonHoldt 2012). Based on the recommendations of Janes et al. (2017), we assessed the number of genetic clusters by applying and comparing these two methods. Finally, CLUMPAK software (Kopelman et al. 2015) was used to summarize and visualize STRUCTURE outputs.

### 2.5. Genetic diversity

Single and multilocus estimates of genetic diversity were calculated for each population as the expected heterozygosity (*He*, *sensu* Nei 1978), observed heterozygosity (*Ho*) and the mean number of private alleles (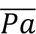) using GenAlEx 6.5 (Peakall & Smouse 2006). In addition, standardized allelic richness (*Ar*) was computed using the rarefaction method of FSTAT. The estimate of deviation from random mating (*F_IS_*) was calculated according to Weir and Cockerham (1984) and statistical significance was computed using GENETIX 4.02 (Belkhir et al. 2004) based on 1,000 permutations.

### 2.6. Statistical analysis

To test the null hypothesis that island and coastline populations did not differ in genetic diversity, a Kruskal-Wallis test was performed for each estimator of genetic diversity (*He*, *Ar* and 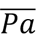), *F_IS_* and *F_ST_* using Minitab (Version 19.2020.2.0). This test was performed on the set of data solely including the 24 continuously sampled populations (S1 to S24). Populations S25 to S32 were excluded as no islands were sampled alongside and because of their high geographic isolation. The same tests were applied to compare each estimator of genetic diversity between *divergent* populations (*i.e.,* associated to the highest *F_ST_* values) and the rest of the populations.

### 2.7. Connectivity model

#### 2.7.1. Hydrodynamic modelling

Flow fields were provided by a 3D regional configuration (MANGA2500) of a hydrodynamic model (MARS3D, Petton et al. 2023). The model domain ranges from 41°N to 55°N in latitude, and 18°W to 9°30′E in longitude, encompassing all studied populations from southern Brittany to the Strait of Dover. The domain is resolved horizontally by a regular 2.5 km grid and vertically by 40 sigma levels. The model was forced by meteorological conditions obtained from the Météo-France ARPEGE model and AROME for the nested zooms with spatial and temporal resolutions of 2.5 km and 1 h, respectively. At the boundaries of the global 3D domain, the forcing by tides was provided by a larger 2D model covering the northwest Atlantic, forced by 14 tidal constituents (Lyard et al. 2006).

#### 2.7.2. Lagrangian Modelling

The Lagrangian trajectory of the particles (meiospores or fertile parts of the thallus) was simulated using ICHTHYOP v.3.3.6 (Lett et al. 2008), which employed the hourly 3D flow fields as input data. Lagrangian simulations were performed from June to October, corresponding to the period with the highest peak in the fertility of *L. digitata* (Bartsch et al. 2008). Particles were released within circles with a radius of 2 km and were allowed to drift for 15 days to account for dispersal by fertile thallus and by meiospores while reducing the computational time of simulations. The computational time step of the Lagrangian model was set to 300s and the position of the particle was recorded every 20 min. For each simulation, 5,000 particles were released every two hours. This configuration was applied for each location and repeated for eight days per month using the hydrodynamic outputs of years 2014 and 2015. The eight days of simulation were chosen by taking two days per week (chosen randomly) in order to consider tidal variation during a month while reducing computational time. A total of 960 dispersal events were hence simulated over two years for each site. Pairwise oceanographic connectivity (*P_ij_*) was estimated following Reynes et al. (2021) and is defined as: *P_ij_* = *N_i-j_*/*N_i_*, with *N_i_* corresponding to the total number of released particles from site *i* and *N_i-j_* the total number of released particles from *i* arriving at *j* within 15 days of dispersal. *P_ij_* was then averaged over the full period of the simulation to obtain the mean connectivity matrix. The level of self-recruitment (*P_ii_*) was estimated with the same method as *P_ij_* but using a threshold of 48h to distinguish emitted particles (from the release area) from those returning to the release area after 48h.

### 2.8. Geographic distance and habitat discontinuity

Using ArcGIS 10, we first extracted the bathymetric contour from a 0.001°-resolution digital terrain model (or DTM, data.shom.fr). Given the shape of the peninsula of Brittany, and in order to standardize the calculation of pair-wise distances between coastal populations, distances were calculated following the 5m isobath. When at least one of the populations was situated on an island, the straight-line distance across the sea to another island or to the closest coastline population was considered. A 0.009°-resolution raster layer representing the distribution of rocky seabed in the English Channel and around Brittany was prepared from the best available substrate information available at Ifremer in May 2019. The information on the spatial distribution of bedrock comes from the combination of several sources: 1) maps of coastal habitats produced as part of the Rebent project, with the scales varying from 1/2,000 to 1/10,000; 2) maps of existing habitat produced as part of the MESH project, with the scales varying from 1/50,000 to 1/10,0000 and 3) from the local extraction of the information on the presence of rock from topobathymetric Lidar data (DTM of 5 m resolution) (*e.g*., Bajjouk et al. 2015). This layer was subsequently used to compute the continuity of rocky substrata between pairs of sites. Only the rock extents situated above 5m depth (corresponding to 5m below the lowest astronomical tide) were considered in this calculation given that *L. digitata* is considered unable to colonize areas below this depth (Robuchon et al. 2014, Figure 1). The proportion of the geographic distance unoccupied by rocky substrata was considered for subsequent analyses to prevent correlation with geographic distance.

### 2.9. Temperature data

Sea-surface temperature (SST) were derived from daily mean satellite imagery (Copernicus product ID: SST_GLO_SST_L4_REP_OBSERVATIONS_010_011) for the years 2016 and 2017 using 0.05°-resolution data obtained in 2019 from the EU Copernicus Marine Service (see Good et al., 2020 for product details). The minimum, maximum, average and range of annual SST were calculated for each population.

### 2.10. Linking environmental variables to genetic structure

Global and partial distance-based redundancy analyses (db-RDA, Legendre and Andersson 1999) were conducted to investigate the individual and joint effects of geographic distance, dispersal mediated by ocean currents, sampling years, SST (minimum, maximum, mean and range) and habitat discontinuity on the total explained genetic variation. The analyses were repeated for domains of different size to assess if and how the contribution of each predictor varied according to it. The “overall scale” corresponded to our whole dataset (*i.e.*, 32 populations), the “regional scale” accounted for the 29 populations from Brittany (S1 to S29) and a “finer scale” included solely the 24 continuously sampled populations (S1 to S24). Sampling years were taken into account to assess how variation in sampling year explained the contemporary genetic differentiation. The first two axes of the PCoA performed on the *F_ST_* matrix (see “Population Structure” subsection above) were used as the response variables representing the core of the genetic structure in the db-RDA. The environmental explanatory variables accounting for geographic distance and ocean currents were respectively transformed into distance-based Moran’s eigenvector maps (or dbMEM, Dray et al. 2006) and Asymmetric Eigenvector Maps (or AEM, Blanchet et al. 2008). dbMEMs are derived from spectral graph theory and permit describing the spatial autocorrelation between sampling locations using orthogonal variables (corresponding to eigenfunctions, Dray et al. 2006). dbMEM eigenvectors result from the matrix of geographic distance and were computed with the “adespatial” package using the default settings for truncations. Only the positive eigenvalues were retained as no negative autocorrelation between sampling positions was expected (Dray et al. 2006). The construction of AEMs uses the same framework while accounting for asymmetric directional spatial processes (Blanchet et al. 2008). The mean oceanographic connectivity matrix (see “Lagrangian Modeling” subsection) was transformed into a nodes-to-edges matrix and the edges were weighted by the probability of dispersal before calculating AEM eigenvectors with the “adespatial” package.

The calculations of dbMEM and AEM eigenvectors were iterated at each domain size following the aforementioned method. It is worth noting that the first eigenvectors in both dbMEMs and AEMs (*e.g*., AEM1, dbMEM1) are associated to broad-scale patterns while the highest eigenvectors (e.g., dbMEM5, AEM25) highlight fine-scale patterns.

For the global db-RDA, an elastic net regularized regression was performed using the “glmnet” package (Friedman et al. 2010) with an elastic net mixing parameter α fixed at 0.08 to facilitate the identification of irrelevant variables and the highly correlated ones. This method is considered successful when the number of explanatory variables exceeds the sample size (Ogutu et al. 2012). The remaining variables were then selected via a stepwise forward procedure using the ordiR2step function from the “vegan” package (Oksanen et al. 2013). This function selects variables to build the *optimal* model, defined as the model maximizing the adjusted coefficient of determination (R^2^), while minimizing the *p*-value (Blanchet et al. 2008). Finally, an analysis of variance (ANOVAs; 1,000 permutations) was performed to assess the significance of the model, axes and retained variables.

## 3. RESULTS

### 3.1. Prior analyses

The PCA run on the whole dataset allowed to identify 12 *L. hyperborea* individuals (Figure S1, Table S1), a sister species to *L. digitata* with an overlapping range distribution (Robuchon et al. 2014). Their identities were additionally confirmed based on allelic size at markers Ld148, Ld158, Ld704, Lo4-24 and Lo454-23 following Mauger et al. (2021). These individuals were sampled in populations S6 (2 ind.), S9 (1 ind.), S21 (3 ind.), S22 (3 ind.), S25 (1 ind.), S28 (1 ind.) and S30 (1 ind.) and were removed from our dataset, giving a total of 896 individuals. Null alleles were present in several populations (Table S2). However, differences between F_ST_ values in the pairwise comparison were of order 10^−3^ (data not shown). Therefore, we concluded that the frequency of null alleles was negligible and our dataset was analyzed without taking into account correction for null alleles. In addition, the correlation between the number of alleles and the number of genotypes with missing data (*p*-value = 0.462) suggest that the presence of null alleles is negligible. Finally, no significant genotypic linkage disequilibrium was obtained, neither within nor across populations (Table S3).

### 3.2. Population Structure

According to both ΔK and ln Pr(X|K) methods, K = 2 appears to be the optimal value for genetic clusters (Figure S2). The barplot for K = 2 illustrates a gradual north-wise differentiation, with southern Brittany (S1-S4, referred to as “SB” hereafter) and the Bay of Saint-Malo (S26-S29, referred to as “SMB” hereafter) being particularly well separated (Figure 2A). ΔK method suggest K = 3 to be the next optimal value, followed by K = 12, while the ln Pr(X|K) method first indicates K = 12 and then K = 3 (Figure S2). The barplot for K = 3 separates SB in another cluster (Figure 2B). The one for K = 12 clearly separates SB, southwest Brittany (S5-S9), Locquirec (S21), Roche-Douvres (S25), SMB, Normandy (S30-S31) and Wissant (S32), while western and northern populations (S10-S20, S22-S24) are highly admixed (Figure 2C). This barplot therefore indicates eight main genetic groups: SB (S1-S4), southwest Brittany (S5-S9), Locquirec (S21), populations from western and northern Brittany (S10-S20, S22-S24), Roche-Douvres (S25), SMB (S26-S29), Normandy (S30-S31) and Wissant (S32). Among the eight genetic groups, populations from SB, Locquirec, SMB and Wissant were associated to significantly higher mean *F_ST_* values (Table 2, and Table S4 for pairwise *F_ST_* values and their significance; ANOVA, *p*-value = 5.12e^−11^). From now on, these ten populations will be termed as *divergent* populations. The high mean *F_ST_* obtained for S32 can be interpreted from allele frequencies spectra, in particular at loci Ld148, Ld371 and Lo454-17. However, allelic spectra were less informative for the remaining populations. Although STRUCTURE results suggest that island populations are generally well admixed with coastline populations except for Roche-Douvres (S25, Figure 2C), the *F_ST_* values among coastline populations were significantly lower than among islands or between coastline and island populations (Kruskal-Wallis test, *p*-value = 0.034).

**Table 2.** Mean *F_ST_* over all pairwise comparisons are given for the group of populations that appeared to be particularly differentiated. S1-S4 corresponds to the southernmost populations (SB); S21 to Locquirec; S26-S29 to populations from Saint-Malo Bay (SMB), S32 to the population from Dover Strait, and “Others” correspond to the rest of the populations. Minimum and maximum *F_ST_* values between pairs of sites are also indicated.

| Populations | Mean $F_{ST}$ | $F_{ST}$ min | $F_{ST}$ max |
| --- | --- | --- | --- |
| S1-S4 | 0.183 | 0.024 | 0.395 |
| S21 | 0.168 | 0.107 | 0.317 |
| S26-S29 | 0.144 | 0.019 | 0.395 |
| S32 | 0.227 | 0.159 | 0.356 |
| Others | 0.08 | 0.01 | 0.23 |

### 3.3. Genetic diversity

Estimates of genetic diversity averaged over the 11 markers are provided in Table 3. Most quantities varied by a factor of two to three across populations: the lowest value was always associated to S32 and the highest to populations located on Brittany’s western-most sites (S8-S14, Table 3). Variation in genetic diversity across populations was the highest for allelic richness, with a minimum value of 2.542 (S32) and a maximum of 6.151 (S12, Table 3, Figure 3). Departure from HW equilibrium (*F_IS_*) showed a significant deficit of heterozygotes for several populations (Table 3) and repeated Multilocus genotypes (MLG) were detected in S4 (1 MLG in 2 individuals) and S28 (1 MLG in 2 individuals). The presence of MLG was not correlated with any significant deviation from HW equilibrium (Table 3).

**Figure 3.**
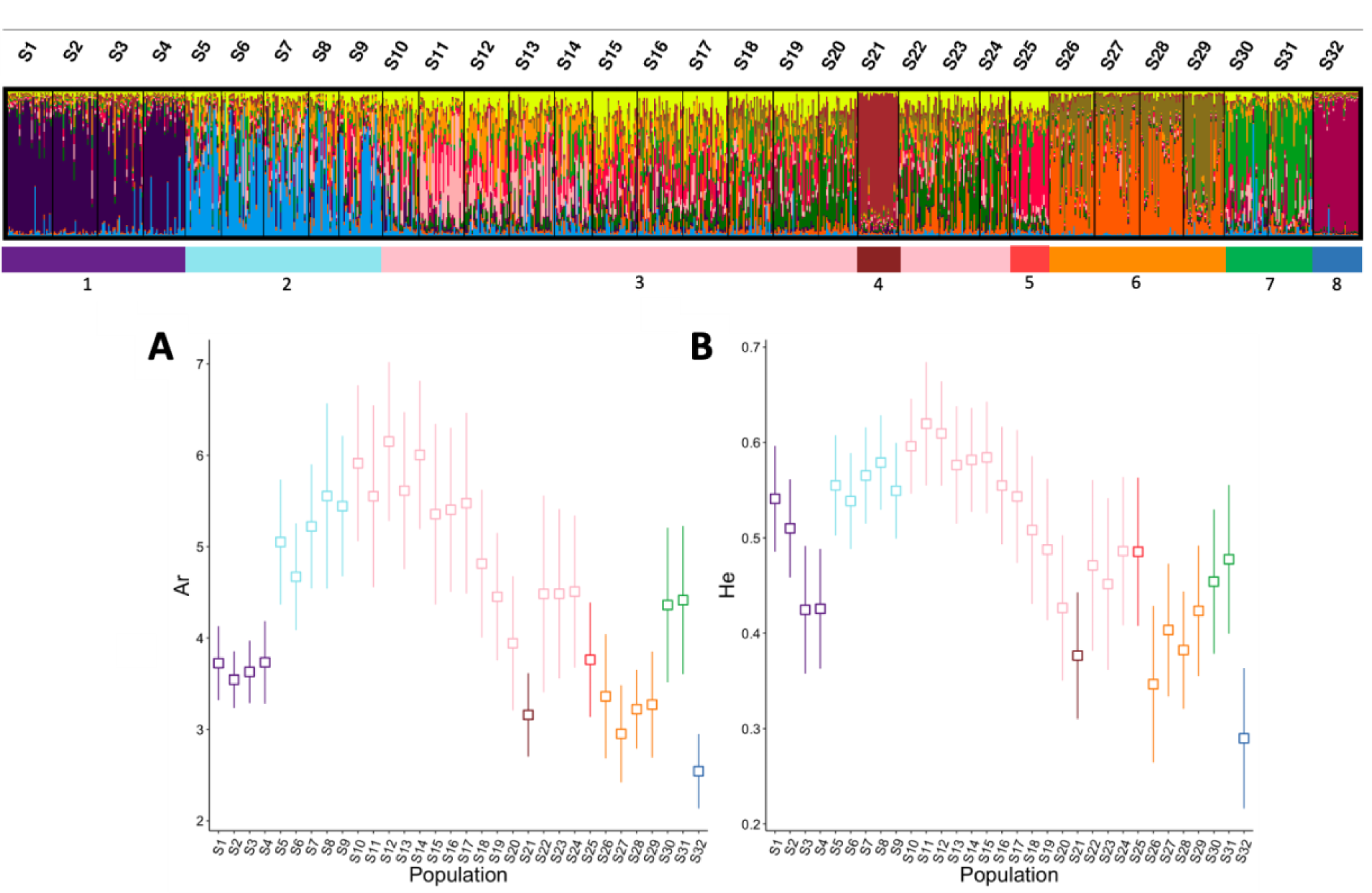
Comparison of **A.** allelic richness (*Ar*) and **B.** expected heterozygosity (*He*) across space. The colors of the plots correspond to the eight genetic groups which are illustrated with the STRUCTURE barplot above the figures. These figures give the average value for each of the 32 populations of our study, the standard deviation observed across markers is illustrated.

**Table 3.**
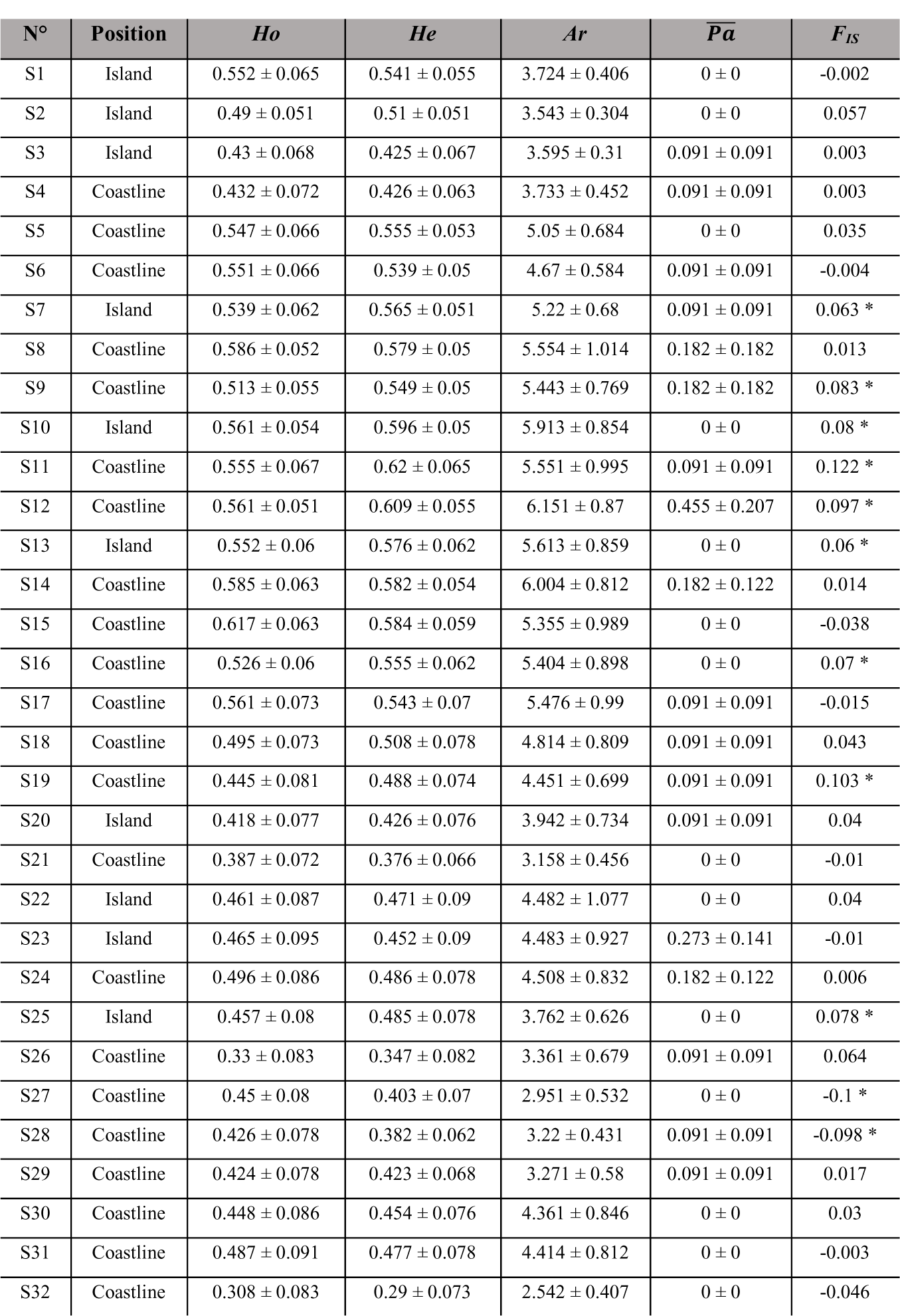
Estimates of genetic diversity associated to each population using *Ho*: observed heterozygosity, *He*: expected heterozygosity, *Ar*: allelic richness (calculated with minimum 19 diploid individuals), 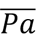: mean private alleles and deviation from Hardy-Weinberg Equilibrium estimated using *F_IS_*, * when the deviation is significant (*p*-value < 0.001, based on 1,000 permutations using GENETIX).

| N° | Position | *Ho* | *He* | *Ar* | $\overline{P\alpha}$ | $F_{IS}$ |
| --- | --- | --- | --- | --- | --- | --- |
| S1 | Island | 0.552 ± 0.065 | 0.541 ± 0.055 | 3.724 ± 0.406 | 0 ± 0 | -0.002 |
| S2 | Island | 0.49 ± 0.051 | 0.51 ± 0.051 | 3.543 ± 0.304 | 0 ± 0 | 0.057 |
| S3 | Island | 0.43 ± 0.068 | 0.425 ± 0.067 | 3.595 ± 0.31 | 0.091 ± 0.091 | 0.003 |
| S4 | Coastline | 0.432 ± 0.072 | 0.426 ± 0.063 | 3.733 ± 0.452 | 0.091 ± 0.091 | 0.003 |
| S5 | Coastline | 0.547 ± 0.066 | 0.555 ± 0.053 | 5.05 ± 0.684 | 0 ± 0 | 0.035 |
| S6 | Coastline | 0.551 ± 0.066 | 0.539 ± 0.05 | 4.67 ± 0.584 | 0.091 ± 0.091 | -0.004 |
| S7 | Island | 0.539 ± 0.062 | 0.565 ± 0.051 | 5.22 ± 0.68 | 0.091 ± 0.091 | 0.063 * |
| S8 | Coastline | 0.586 ± 0.052 | 0.579 ± 0.05 | 5.554 ± 1.014 | 0.182 ± 0.182 | 0.013 |
| S9 | Coastline | 0.513 ± 0.055 | 0.549 ± 0.05 | 5.443 ± 0.769 | 0.182 ± 0.182 | 0.083 * |
| S10 | Island | 0.561 ± 0.054 | 0.596 ± 0.05 | 5.913 ± 0.854 | 0 ± 0 | 0.08 * |
| S11 | Coastline | 0.555 ± 0.067 | 0.62 ± 0.065 | 5.551 ± 0.995 | 0.091 ± 0.091 | 0.122 * |
| S12 | Coastline | 0.561 ± 0.051 | 0.609 ± 0.055 | 6.151 ± 0.87 | 0.455 ± 0.207 | 0.097 * |
| S13 | Island | 0.552 ± 0.06 | 0.576 ± 0.062 | 5.613 ± 0.859 | 0 ± 0 | 0.06 * |
| S14 | Coastline | 0.585 ± 0.063 | 0.582 ± 0.054 | 6.004 ± 0.812 | 0.182 ± 0.122 | 0.014 |
| S15 | Coastline | 0.617 ± 0.063 | 0.584 ± 0.059 | 5.355 ± 0.989 | 0 ± 0 | -0.038 |
| S16 | Coastline | 0.526 ± 0.06 | 0.555 ± 0.062 | 5.404 ± 0.898 | 0 ± 0 | 0.07 * |
| S17 | Coastline | 0.561 ± 0.073 | 0.543 ± 0.07 | 5.476 ± 0.99 | 0.091 ± 0.091 | -0.015 |
| S18 | Coastline | 0.495 ± 0.073 | 0.508 ± 0.078 | 4.814 ± 0.809 | 0.091 ± 0.091 | 0.043 |
| S19 | Coastline | 0.445 ± 0.081 | 0.488 ± 0.074 | 4.451 ± 0.699 | 0.091 ± 0.091 | 0.103 * |
| S20 | Island | 0.418 ± 0.077 | 0.426 ± 0.076 | 3.942 ± 0.734 | 0.091 ± 0.091 | 0.04 |
| S21 | Coastline | 0.387 ± 0.072 | 0.376 ± 0.066 | 3.158 ± 0.456 | 0 ± 0 | -0.01 |
| S22 | Island | 0.461 ± 0.087 | 0.471 ± 0.09 | 4.482 ± 1.077 | 0 ± 0 | 0.04 |
| S23 | Island | 0.465 ± 0.095 | 0.452 ± 0.09 | 4.483 ± 0.927 | 0.273 ± 0.141 | -0.01 |
| S24 | Coastline | 0.496 ± 0.086 | 0.486 ± 0.078 | 4.508 ± 0.832 | 0.182 ± 0.122 | 0.006 |
| S25 | Island | 0.457 ± 0.08 | 0.485 ± 0.078 | 3.762 ± 0.626 | 0 ± 0 | 0.078 * |
| S26 | Coastline | 0.33 ± 0.083 | 0.347 ± 0.082 | 3.361 ± 0.679 | 0.091 ± 0.091 | 0.064 |
| S27 | Coastline | 0.45 ± 0.08 | 0.403 ± 0.07 | 2.951 ± 0.532 | 0 ± 0 | -0.1 * |
| S28 | Coastline | 0.426 ± 0.078 | 0.382 ± 0.062 | 3.22 ± 0.431 | 0.091 ± 0.091 | -0.098 * |
| S29 | Coastline | 0.424 ± 0.078 | 0.423 ± 0.068 | 3.271 ± 0.58 | 0.091 ± 0.091 | 0.017 |
| S30 | Coastline | 0.448 ± 0.086 | 0.454 ± 0.076 | 4.361 ± 0.846 | 0 ± 0 | 0.03 |
| S31 | Coastline | 0.487 ± 0.091 | 0.477 ± 0.078 | 4.414 ± 0.812 | 0 ± 0 | -0.003 |
| S32 | Coastline | 0.308 ± 0.083 | 0.29 ± 0.073 | 2.542 ± 0.407 | 0 ± 0 | -0.046 |

The ten divergent populations were associated with the lowest level of genetic diversity (Kruskal-Wallis tests, *p*-value for *He:* < 1.0e^-4^, *Ar*: < 1.0e^-4^, see Figure 3) although the same tendency was not observed for 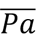 (Kruskal-Wallis test, *p*-value: 0.274). These populations also appear to have significantly lower *F_IS_* values compared to the rest of the populations (Kruskal-Wallis test, *p*-value = 0.022). In contrast, while genetic diversity estimates were always higher in coastline than in island populations, differences were not significant based on Kruskal-Wallis tests (*p*-value for *He*: 0.212, *Ar*: 0.108 and 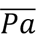: 0.099).

Similarly, higher *F_IS_* values were observed in island populations but the difference was, again, not significant (Kruskal-Wallis tests, *p*-value: 0.739).

### 3.4. Habitat discontinuity

The proportions of habitat discontinuity shown in Table S5 suggest a patchy pattern with several populations that appeared particularly isolated. As illustrated by Figure 1, island populations appear naturally isolated by the absence of continuous favorable substrate (*i.e*., rocky substrata above 5m depth) connecting them to the nearest mainland or island populations. Populations from SB and the north of the Cotentin Peninsula (S30-S32), appeared highly isolated from the rest of the populations but also within themselves thus revealing high habitat fragmentation. At the same time, the expanse of rocky habitat appears continuous along the northern Brittany coast (S15-S29) as shown by the small proportion of geographic distance which is unoccupied by rocky substrata (Table S5). The increase in habitat discontinuity between southern (< S11) and northern populations (> S11) was mainly driven by the topography of the Rade of Brest and its lack of rocky substrata at the 5 m isobath, and even above.

### 3.5. Oceanographic connectivity

Connectivity probabilities resulting from Lagrangian simulations are shown in Table S6. In most cases, the highest probabilities of connectivity were associated with neighboring populations. Contrastingly, groups of populations such as SB, SMB and Normandy appeared highly connected within the group, but largely unrelated to other populations (Table S6). The probability of connectivity of S32 with any other population was null, highlighting its strong isolation. In the case of S21, while a particle released from this population attains neighboring ones, particles released from the latter had very slight probabilities to recruit in S21 (Table S6). The difference between divergent and non-divergent populations was well marked when comparing the number of connectivity links, defined as the mean number of populations with which the probability of connectivity is non null. Indeed, spores released from divergent populations reach an average of 5.2 populations compared to 7.9 for non-divergent populations (Table S6). Similarly, spores recruiting in divergent populations originated from an average of 4.9 populations, compared to 9.1 for non-divergent ones (Table S6). The difference in connectivity was less pronounced between island and coastal populations: spores released from islands reach on average 8.5 other populations, compared to 6.6 for coastal populations (Table S6) and spores that recruited in islands originated from an average of 9.2 populations, compared to 6.5 for coastal populations (Table S6).

In terms of directionality, dispersal tends to be oriented in a north-western direction among populations on the south of the Armorican Peninsula (S1-S7) and towards a north-eastern direction when considering north-western and northern populations (S10-S25, Table S6): for instance, a spore released from S10 reaches S11 to S14 with a higher probability than in the opposite direction (S7-S9, Figure 4, Table S6). However, the pattern becomes less clear in the SMB populations (S26-S32). Indeed, as previously stated, these populations are highly connected within each genetic group (SMB and Normandy) but not with populations beyond them, which makes the directionality analysis less relevant.

**Figure 4.**
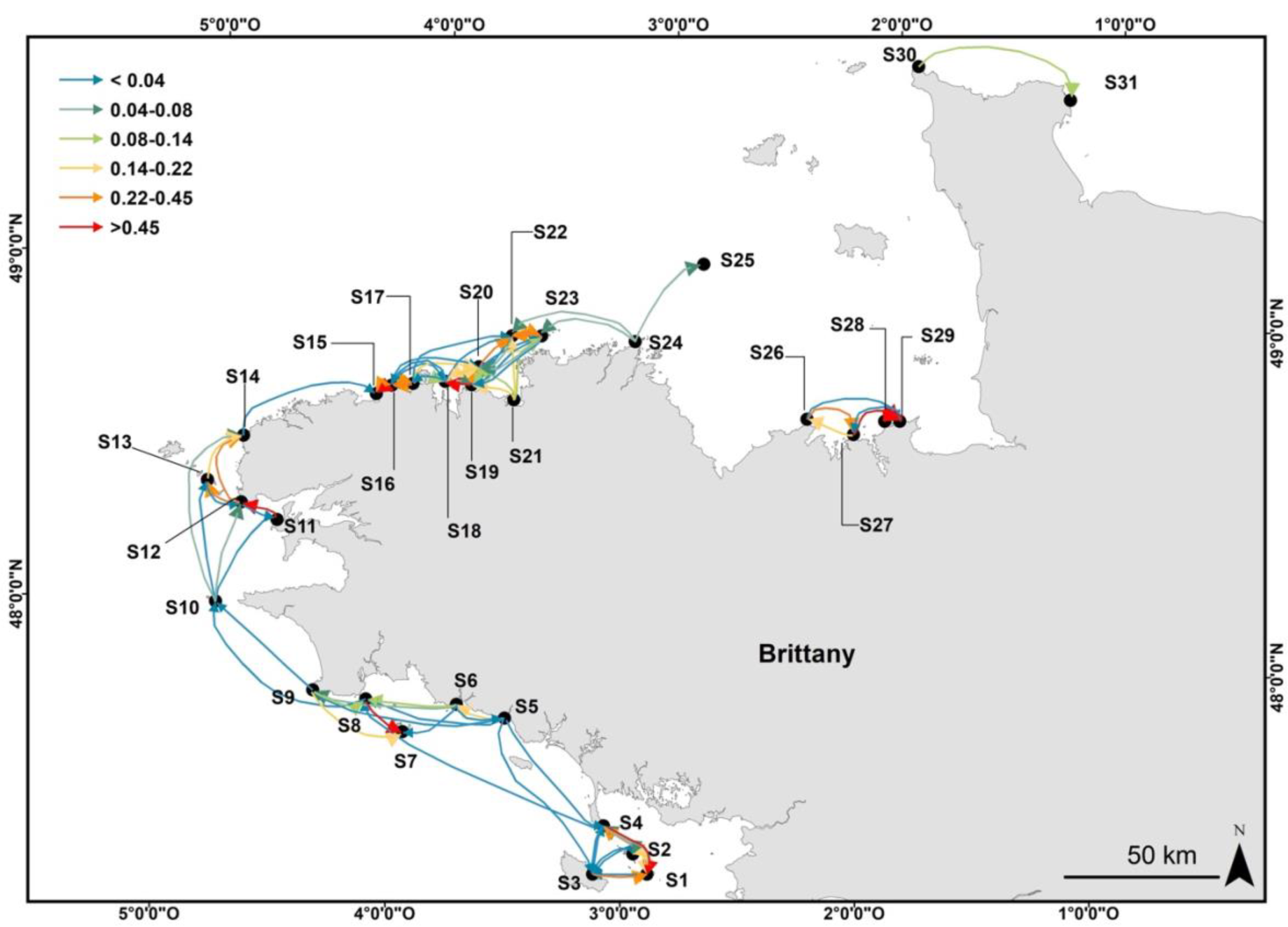
Map illustrating the probability of connectivity obtained from the Lagrangian simulation model according to the color gradient illustrated on the top left corner. Cold color indicates weak connectivity while warm color indicates high connectivity. For the clarity of the figure, values below 10^-2^ are not represented.

On the other hand, Lagrangian simulations provided numbers on self-recruitment probability, which varied from 0.005 (S8) to 0.476 (S25, Table S6). This probability did not significantly differ between island and coastal populations (0.211 and 0.189, respectively, with an ANOVA *p*-value = 0.068), nor between divergent and non-divergent populations (0.255 and 0.169, respectively, with an ANOVA *p*-value = 0.695).

### 3.6. Hierarchical effects of environmental factors on genetic structure

Whichever the geographical scale, habitat discontinuity and sampling years were not retained in the global db-RDA, while the partial db-RDA accounting for these two variables individually were not significant (Table 4). Therefore, for the sake of parsimony, these two variables will not be mentioned in the following results.

**Table 4.**
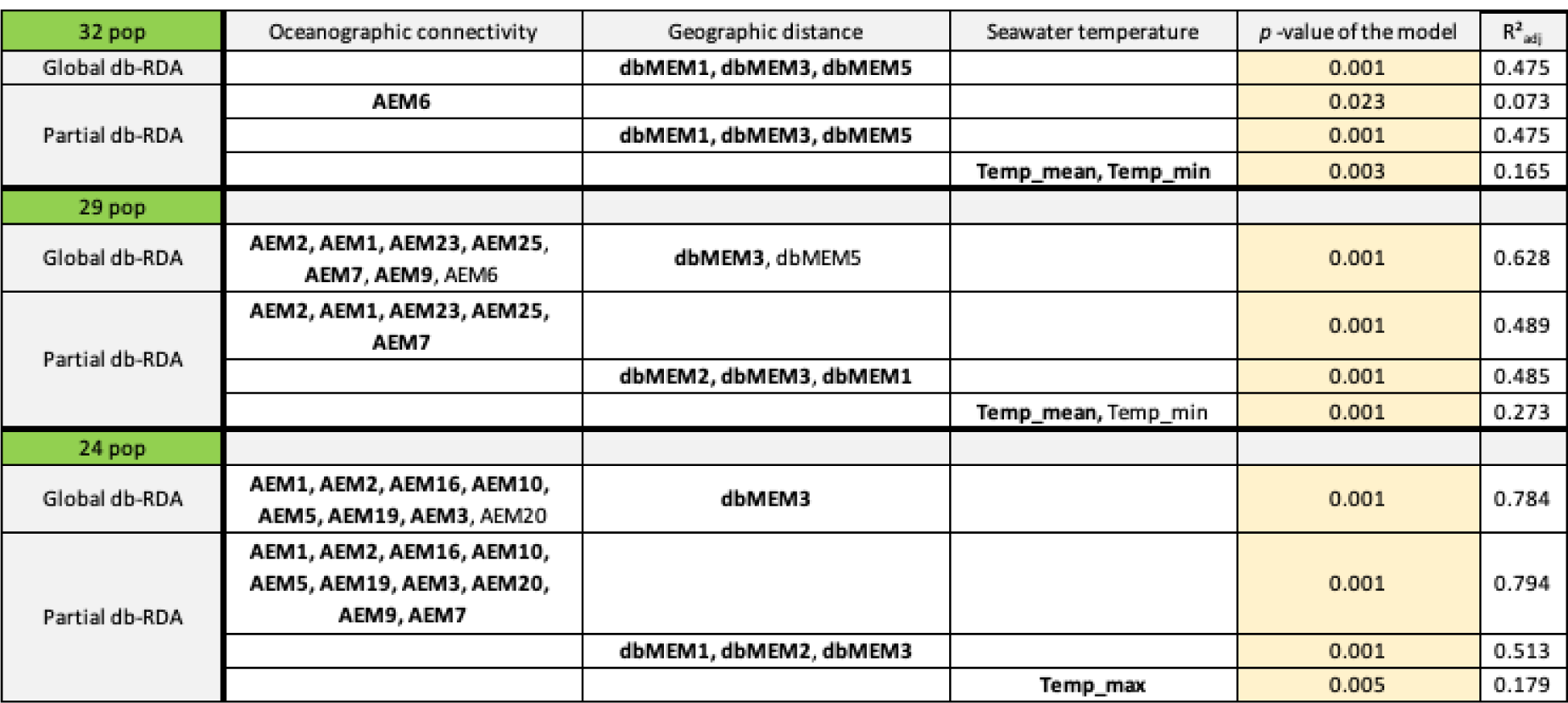
Results of the partial and global db-RDA for each spatial scale (32, 29 and 24 populations). Environmental predictors selected by the stepwise forward selection (ordiR2step) are included in the db-RDA framework, the predictors highlighted in bold are significant at *p* < 0.001 using ANOVA. The adjusted coefficient of determination (R^2^) and the *p*-value of the model are reported. Habitat continuity and sampling year were not selected as a significant predictor in the global nor partial db-RDA and are therefore not represented for a matter of clarity.

#### 3.6.1. Overall scale (32 populations)

The model and the two first axes associated to the global db-RDA were significant (*p*-value < 0.001) with an adjusted coefficient of determination (R^2^) of 0.475 (Table 4). Only three variables were selected by the ordiR2step function which were all related to geographic distance (dbMEM1, 3 and 5, Table 4, Figure 5), suggesting that the latter factor is the main driver of genetic structure at this scale. The first axis, accounting for 78.8% of the variance clearly separates SB from SMB. The second axis, accounting for 17.8% of the variance, mostly isolates S32 whilst populations from Normandy remain poorly differentiated (Figure 5). When partitioning the respective effects of oceanographic connectivity, geographical distance and SST using partial db-RDA, each predictor appeared significant. Geographic distance explained the greatest amount of variance with R^2^ = 0.475, against R^2^ = 0.165 for SST (minimum and mean) and 0.073 for oceanographic connectivity (Table 4).

**Figure 5.**
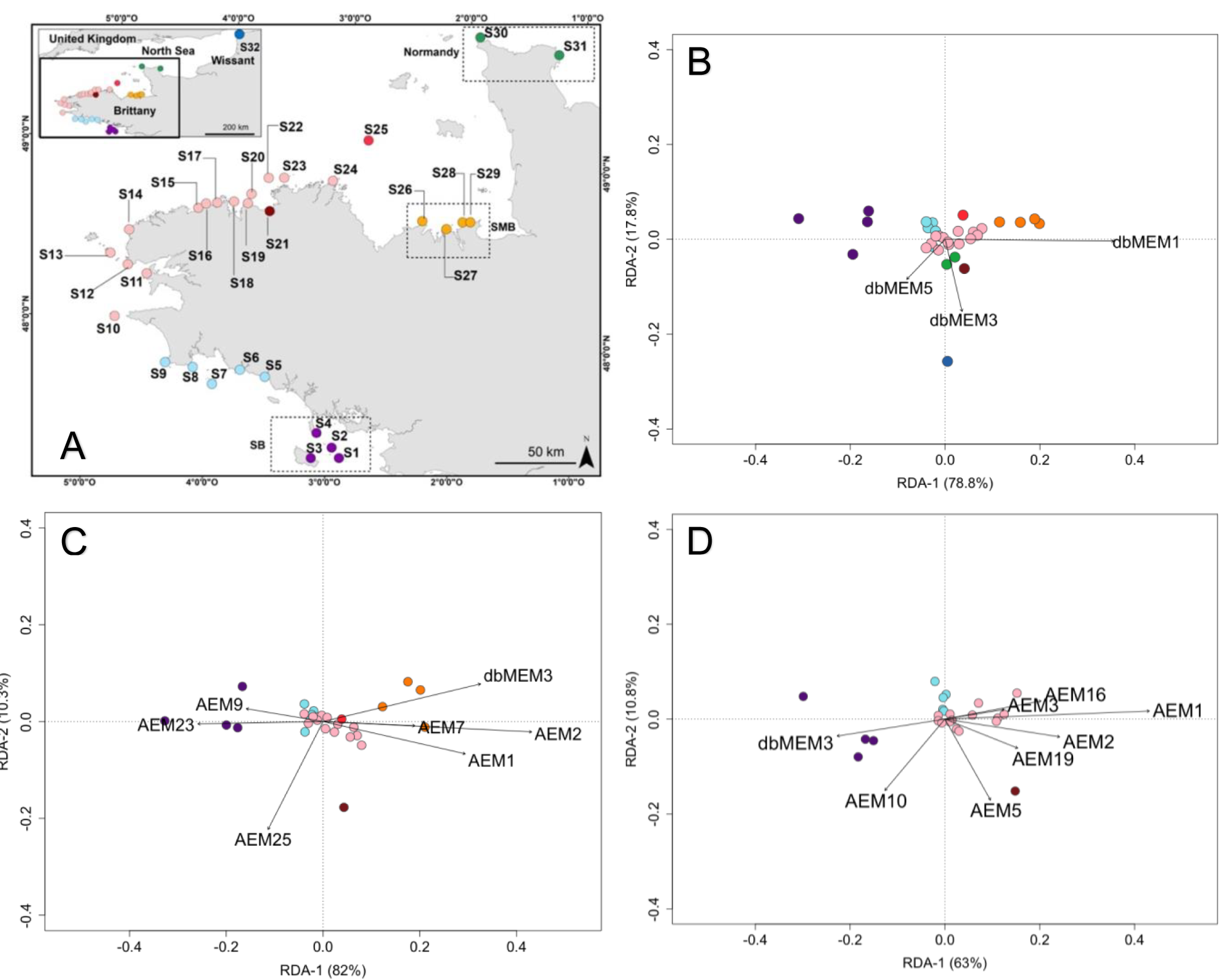
Figures **B**. **C** and **D** represent the results from the global db-RDA ran on B. 32 populations, C. 29 populations and D. 24 populations. The color used for the populations corresponds to the eight genetic clusters as illustrated in Figure 3. The numbers written in brackets on each axis correspond to the percentage of variance explained by each axis. The black arrows illustrate the relative contribution of the significant environmental factors obtained by ANOVA and the stepwise forward selection (ordiR2step) process. The length of the arrows illustrates the elative contribution of each environmental predictor: as the length increases, the relative contribution of the environmental predictor to predict the neutral genetic variation increases.

#### 3.6.2. Brittany scale (29 populations)

To investigate the relative contribution of each environmental factor at a finer scale, populations S30, S31 and S32 were excluded because of their high geographic isolation. The model and the two first axes associated to the global db-RDA were again significant (*p*-value < 0.001) with R^2^ = 0.628 (Table 4). Nine variables were retained by the ordiR2step selection process: two account for geographic distance (dbMEM3 and 5, Table 4) and seven for oceanographic connectivity (AEM1, 2, 6, 7, 9, 23 and 25, Table 4). The first axis (82% of the variance) again clearly separates populations according to their latitude and populations from the SMB appear isolated due to geographic distance. The second axis (10.3%) is mainly driven by AEM25 and isolates S21, suggesting the role of oceanographic currents on its genetic isolation (Figure 5). When partitioning the respective effects of oceanographic connectivity, geographical distance and SST, each predictor appeared significant. At this scale, oceanographic connectivity and geographic distance explained approximately the same amount of variance (R^2^_adj_ = 0.489 and 0.485, respectively) while R^2^ = 0.273 for SST (mean, Table 4). Overall, this indicated that excluding the most isolated populations (farther than 600 km from the rest of the populations) accentuated the effects of dispersal mediated by ocean currents.

#### 3.6.3. Continuous scale (24 populations)

A third analysis accounting for the 24 continuously distributed populations (S1-S24) was performed by additionally excluding SMB populations. At this scale, the highest distance between two neighboring populations is reduced to ca. 160 km (between S11 and S12 when following coastline, Figure 4) and no major gap in connectivity was reported (Figure 4). The model and the three first axes were significant (*p*-value < 0.001 for the model and the first axis, *p*-value < 0.05 for the remaining two axes) with R^2^ = 0.777 (Table 4). Nine variables remained after the two selection processes: eight were associated to oceanographic currents (AEM1, 2, 3, 5, 10, 16, 19 and 20) and one to geographic distance (dbMEM3, Table 4). At this scale, oceanographic connectivity explained a greater amount of variance than geographic distance (R^2^_adj_ = 0.794 and 0.513, respectively), while SST explained 17.9% of the variance (maximum SST). It is worth noting that at this scale, the genetic differentiation between Southern (S1-S9) and Northern Brittany (S10-S25) is widely explained by oceanographic processes (Figure 5). This first axis is mostly explained by AEM1, denoting the gradual north-wise differentiation, while AEMs of higher order mainly indicate the strong density of connectivity links among northern populations (S10-S25). The second axis again isolates S21, which is mostly explained by AEM5. Overall, the amount of variation explained by seascape features increases as the spatial scale is reduced, ranging from R^2^ values of 0.475 to 0.628 and 0.777 at this semi-continuous scale.

## 4. DISCUSSION

Our study builds on previous microsatellite research (Billot et al. 2003, Valero et al. 2011, Couceiro et al. 2013, Robuchon et al. 2014) to refine our understanding of *Laminaria digitata*’s population structure through a comprehensive sampling design on the southern edge of its East Atlantic distribution. We applied a distance-based redundancy analysis (db-RDA) to disentangle the relative contribution of habitat discontinuity, geographical distance, hydrodynamic processes, sea surface temperature (SST) and sampling year on the observed genetic differentiation. Unlike previous studies carried on *L. digitata* (Billot et al., 2003; Valero et al., 2011) and on other kelp species (Alberto et al. 2010, 2011, Selkoe et al. 2010, Durrant et al. 2018), our results indicated a limited effect of habitat fragmentation compared to the contribution of ocean currents and geographic distance. Moreover, we found that the relative effect of these two latter environmental variables varied according to the extent of the considered domain: while geographic isolation was more relevant at our ‘overall scale’ (> 600 km), oceanographic processes were the main drivers of genetic structure at a finer spatial resolution (< 300 km). We will now discuss these different results in the light of their possible impact on conservation policies applied to this species at its southern range limit.

### Mosaic patterns of well-connected and divergent populations

Previous results on *L. digitata* have shown significant population differentiation from 1 to 10 km (Billot et al. 2003, Valero et al. 2011, Robuchon et al. 2014), a fine-scale genetic structure supported by the commonly reported short dispersal distance in kelps (Dayton 1985, Norton 1992). Contrastingly, our results suggest that genetic differentiation was not significant between several pairs of populations separated by more than 50 km, particularly for populations located in western and north-western Brittany. In fact, the discrete sampling schemes employed in those previous studies might have led to the description of discrete population structure, yet our results pointed out how sampling scheme variation can lead to different conclusions in this species (see also Serre & Pääbo 2004 and Bradburd et al. 2018). The same bias might affect the results obtained in the Bay of Saint-Malo (SMB, S26-S29) and Normandy, which could be improved in further studies by increasing sampling sites to avoid any gaps (*e.g.*, encompassing populations from Jersey and Guernsey). Nonetheless, finer sampling scheme might not modify the results obtained for Wissant (S32) given the absence of nearby populations (Araújo et al. 2016), an observation which is concordant with the fact that this species has been classified as “completely loss” in this region (de Bettignies et al. 2021). The extinction of *L. digitata* from the Pas-de-Calais could be predicted by our study given its low values of neutral diversity and its lack of connectivity which prevents any recovery from potential source populations.

Low genetic differentiation at large geographic distance was also substantial when considering island populations which were separated by more than 100 km from the coast. Although genetic differentiation was significantly higher when considering at least one island population (*i.e*., island-island; island-coastline), both the number of connectivity links (defined as the mean number of populations with which the probability of connectivity is non null) and values of genetic diversity revealed that islands are generally well connected. Nonetheless, this conclusion might be affected by means of dispersal and should not be taken as a general rule. Islands may in fact be less effective in exporting propagules compared to coastline populations in some species (Bell 2008), and this should be acknowledged in fishery management and conservation actions.

Our results enabled to identify genetically divergent populations despite our sampling bias, that were associated with significantly lower values of genetic diversity. This was the case for populations from southern Brittany (SB, S1-S4), Locquirec (S21) and SMB that appeared highly differentiated from surrounding populations at small spatial scale (< 20 km), in addition to the previously mentioned population of Wissant (S32) being particularly impoverished. Our study thus reveals a mosaic of situations with well-connected populations (mainly located in the north-western part of Brittany) surrounded by populations that appeared less connected and genetically less diverse (mainly in the south and east). This pattern is consistent with the ecological and demographic status of *L. digitata* forest which has been described as highly contrasted in the studied aera, ranging from a “no declined reported” status in north-western Brittany to a “local decline” status in southern Brittany and the Bay Saint-Malo (de Bettignies et al. 2021).

Populations from SB and SMB meet the usual expectations of marginal populations (Pironon et al. 2017, Nadeau & Urban 2019), namely, poor genetic variability and high genetic differentiation within clusters. Moreover, repeated multilocus genotypes (MLG) were observed in a population from both SB (S4) and SMB (S28). MLG could result from the fertilization of gametes coming from the same parental gametophytes, which probability should increase as the population size decreases. The presence of MLG would be consistent with low population size in these two populations. These results would therefore corroborate the abundant-center hypothesis (Brown 1984) although finer estimations of population size are required. Another factor that has to be taken into account in the case of SB is that SST in the area reaches 21°C in summer (Gallon et al. 2014), yet sporulation in this species has been shown to be severely impacted at temperatures above 17°C (Bartsch et al. 2013). Although mechanisms linked to sporulation could be adapted to higher temperature, Oppliger et al. (2014) have shown that the mean number of released meiospores is significantly lower in Quiberon (S4) compared to a population in northern Brittany. Yet a decreased amount of released meiospores should lead to a decreased connectivity and thus higher genetic differentiation. This underlines the fact that SST could have various consequences on life history traits in *L. digitata* which should be apparent on data obtained using neutral markers through the phenomenon of isolation by adaptation (Nosil et al. 2009, Schoville et al. 2012). Similarly, the sharp genetic break observed between southwestern populations (S5-S9) and SB could also be attributed to the difference in SST between these two regions as illustrated by Gallon et al. (2014). We therefore conducted a db-RDA to test the effect of SST and other variables that could explain the observed geographical distribution of genetic variation.

### Limited effects of temperature and habitat discontinuity compared to geographic position and oceanic current

The stepwise variable selection procedure applied prior to db-RDA has never selected variables associated with sampling year, SST or habitat discontinuity. Yet SST and habitat discontinuity were found to be significant factors shaping the genetic structure of other kelp species (Alberto et al. 2010, 2011, Selkoe et al. 2010, Johansson et al. 2015). This discrepancy could stem from a limitation to the use of dbMEMs and AEMs, which may underestimate the importance of other environmental variables that show some correlations with one of these Moran’s eigenvector decomposition (Dalongeville et al. 2018). This argument could be particularly valid for SST as the partial db-RDA considering this variable was always significant whichever the domain extent. To overcome this limitation, one could consider hourly temperature data (*e.g.*, using datalogger) which, in addition to its relevance for intertidal species, could decrease the correlation with dbMEMs. One of the technical limitations that could have led to the non-significance of habitat continuity is that our rock layers data come from the combination of various data with different precisions, or due to the fact that it has been constrained by the 5m bathymetric contour. Another argument is that Alberto et al. (2010, 2011) and Selkoe et al. (2010) have measured habitat continuity by looking at kelp coverage rather than proportion of rocky substrata *per se*. Yet, evidence from other kelp species suggests that sporophyte recruitment largely depends on meiospore density ensuring sperm-egg encountering (Reed 1990). By considering kelp coverage rather than rocky substrata, Alberto et al. (2010, 2011) and Selkoe et al. (2010) might have incorporated the effect of meiospores dilution into habitat discontinuity, which was not our case.

Results from db-RDA revealed that populations from SMB appeared genetically isolated due to their geographic position, rather than by tidal gyre occurring in this region (Salomon & Breton 1993) as previously suggested by Billot et al. (2003) and Robuchon et al. (2014). Nonetheless, this result should be again interpreted with caution as the analysis could have been biased by the gap in the sampling scheme between S25 and S26. The db-RDA ran at the smallest scale (24 populations) further separated the contribution of major and minor oceanographic currents. This highlights that ocean currents not only affect long-distance dispersal (*e.g*., between islands and coastline), but also have localized effects as reported for *L. digitata* in Strangford Narrows (Brennan et al. 2014) and for other species of seaweed characterized by low dispersal abilities (Buonomo et al. 2017, Reynes et al. 2021). Therefore, this study did not verify that SST or habitat discontinuity are important drivers of genetic structure contrarily to an impressive part of the literature, especially in kelps and in *L. digitata* as mentioned previously. However, the strong spatial congruence between the effects of habitat fragmentation, SST difference, geographical location and genetic differentiation observed in our study exemplifies the difficulty to disentangle their effects.

The discrepancy between southern and northern Brittany as observed with STRUCTURE analyses was explained by the major currents according to the db-RDA ran at the smallest scale. This discrepancy is in line with the fact that this region corresponds to an oceanographic front: while the main ocean current in the southern coast of Brittany corresponds to the shelf residual current, which has a northwest-ward direction, northern Brittany is dominated by the English Channel residual circulation, which has a northeast-ward direction (Pingree & Le Cann 1989). This genetic break was also reported for other species (Roman & Palumbi 2004, Jolly et al. 2006, Nunes et al. 2021), and also corresponds to the limit between the northern European Sea and the Lusitanian biogeographical provinces (Spalding et al. 2007) which was pointed out to explain the phylogeographic history of this region. If this can also be applied to *L. digitata*, oceanic front might have then contributed to maintain this historical pattern of differentiation. In addition, the directionality observed from the Lagrangian simulations is consistent with the direction of these two major oceanic currents, although directionality may vary throughout the year (Ayata et al. 2010). Directionality has a rather important consequence in regard to the persistence of populations inhabiting the southern and warmest margin (Nadeau & Urban 2019, DuBois et al. 2022). Indeed, if central and marginal populations are facing different environmental conditions, the asymmetry in gene flow can generate some maladaptation in marginal populations, thereby promoting their extinction (Fouqueau & Roze 2021). In fact, a previous common garden experiment has shown that one of the southernmost populations (Quiberon, S4) shows signs of adaptation in the face of an increase in SST compared to a population from Northern Brittany (Roscoff, Liesner et al. 2020). Therefore, north-wise dispersal could benefit the future adaptation of northern populations if temperature increases in a non-latitudinal manner (DuBois et al. 2022).

## Acknowledgements

We thank Eric Thiébaut, Thierry Comtet, Jean-François Arnaud and Ophélie Ronce for helpful discussions and comments. The OSU Pythéas Institute (Marseille) is thanked for the use of the high-performance computing (HPC) cluster. We are also grateful to the Biogenouest genomics core facility (Genomer Plateforme génomique at the Biological Station of Roscoff in particular Gwenn Tanguy) for their technical support. LF was funded the EU project MARFOR Biodiversa/004/2015, Region Bretagne (ARED 2017 REEALG) and the NOMIS foundation. JA is funded by Portuguese national funds through projects UIDB/04326/2020, UIDP/04326/2020, LA/P/0101/2020 and PTDC/BIA-CBI/6515/2020.

